# Schizophrenia, autism spectrum disorders and developmental disorders share specific disruptive coding mutations

**DOI:** 10.1101/2020.09.02.279265

**Authors:** Elliott Rees, Hugo D. J. Creeth, Hai-Gwo Hwu, Wei J. Chen, Ming Tsuang, Stephen J. Glatt, Romain Rey, George Kirov, James T. R. Walters, Peter Holmans, Michael J. Owen, Michael C. O’Donovan

## Abstract

Genes enriched for rare disruptive coding variants in schizophrenia overlap those in which disruptive mutations are associated with neurodevelopmental disorders (NDDs), particularly autism spectrum disorders and intellectual disability. However, it is unclear whether this implicates the same specific variants, or even variants with the same functional effects on shared risk genes. Here, we show that *de novo* mutations in schizophrenia are generally of the same functional category as those that confer risk for NDDs, and that the specific *de novo* mutations in NDDs are enriched in schizophrenia. These findings indicate that, in part, NDDs and schizophrenia have shared molecular aetiology, and therefore likely overlapping pathophysiology. We also observe pleiotropic effects for variants known to be pathogenic for several syndromic developmental disorders, suggesting that schizophrenia should be included among the phenotypes associated with these mutations. Collectively, our findings support the hypothesis that at least some forms of schizophrenia lie within a continuum of neurodevelopmental disorders.

## Introduction

Schizophrenia is a severe psychiatric disorder associated with a decreased life expectancy and marked variation in clinical presentation, course and outcome. The disorder is highly heritable and polygenic, with risk alleles distributed widely across the genome^1^. Common risk alleles collectively contribute to around a third of the genetic liability^2,3^, and at least 8 rare copy number variants have been identified as risk factors^4,5^. Exome-sequencing studies have also shown a contribution to risk from ultra-rare protein-coding variants; the *de novo* mutation rate is modestly elevated above the expected population rate, and there is an excess of ultra-rare damaging coding variants (frequency < 0.0001 in population) in genes with evidence for strong selective constraint against protein-truncating variants (PTVs)^6–9^. *SETD1A* is currently the only gene in which rare coding variants are associated with schizophrenia at genome-wide significance^10^.

All of the rare CNVs known to be associated with schizophrenia also confer risk for NDDs, that is they are pleiotropic^5^. However, these CNVs are, with one exception, multigenic and therefore it is not established that genic pleiotropy exists whereby the same genes within the CNVs increase liability to each of the disorders^11^. The only example of a single-gene schizophrenia susceptibility CNV is *NRXN1*. Consistent with genic pleiotropy, exonic deletions of *NRXN1* increase liability to schizophrenia and NDDs, but there is marked heterogeneity in the exons affected and the deletion sizes^12^, leaving uncertainty as to whether precisely the same mutation can cause schizophrenia and NDD. In contrast, sequencing studies provide strong support for the hypothesis of genic pleiotropy, with genes that are enriched for ultra-rare coding variants in people with NDDs being enriched for *de novo* variants in people with schizophrenia^7,8^. Moreover, *SETD1A* is not only genome-wide significantly associated with schizophrenia, is also is associated with DD^13^

While *SETD1A* is enriched for PTVs in both schizophrenia and DD^10,13^, the degree to which pleiotropic effects across these disorders are confined to the same functional class of variant within genes is unknown. Moreover, although a specific PTV (c.4582-2delAG>-) in *SETD1A* has been observed multiple times in people with schizophrenia and DD^10^, little is known generally about pleiotropic effects from individual rare coding variants in schizophrenia and NDDs (i.e. true or allelic pleiotropy); this is not a trivial point, as only allelic pleiotropy implies that equivalent changes in gene function confer risk to both schizophrenia and NDDs. For example, some PTVs may cause disease by disrupting dosage sensitive genes (i.e. haploinsufficiency), whereas others in the same gene might result in truncated proteins that have pathogenic gain-of-function or dominant-negative effects^14–16^. Moreover, different PTVs in the same gene can affect different transcripts^17^ leading to different effects. Similar considerations apply to missense variants whose functions are usually unknown, but which can have very different functional consequences for the same gene translating to different pathogenic effects^18^.

In the current study, we analysed sequencing data from schizophrenia and new large NDD cohorts to investigate the nature of the pleiotropic effects of rare coding variants on schizophrenia and NDDs. Specifically, given that neurodevelopmental impairment is typically more severe in NDDs than in schizophrenia^19^, we hypothesised that there would be a tendency for pleiotropic genes to be enriched for a more severe class of mutation in NDDs than in schizophrenia. However, in contrast to expectation, we found that genes enriched for specific classes of *de novo* variant in people with NDDs were also enriched for congruent variant classes in people with schizophrenia. We followed this finding with a more stringent, and conservative, test of allelic pleiotropy, and demonstrated an enrichment in schizophrenia of specific variants that have been observed *de novo* in NDDs. Our findings provide strong evidence for true pleiotropic effects from rare coding variants across these disorders, thus indicating that the same changes in gene function can have very different neurodevelopmental and psychiatric outcomes.

## Results

### Genic pleiotropy

In schizophrenia, *de novo* PTVs were significantly enriched in 127 genes associated with NDD through the same mutation class (rate ratio = 4.89; Table 1). No significant enrichment was observed for schizophrenia *de novo* missense variants in PTV NDD genes (Table 1). The excess of *de novo* PTVs in schizophrenia was ∼3.7 fold greater than that observed for missense variants in PTV NDD genes (Table 1).

**Table 1.**
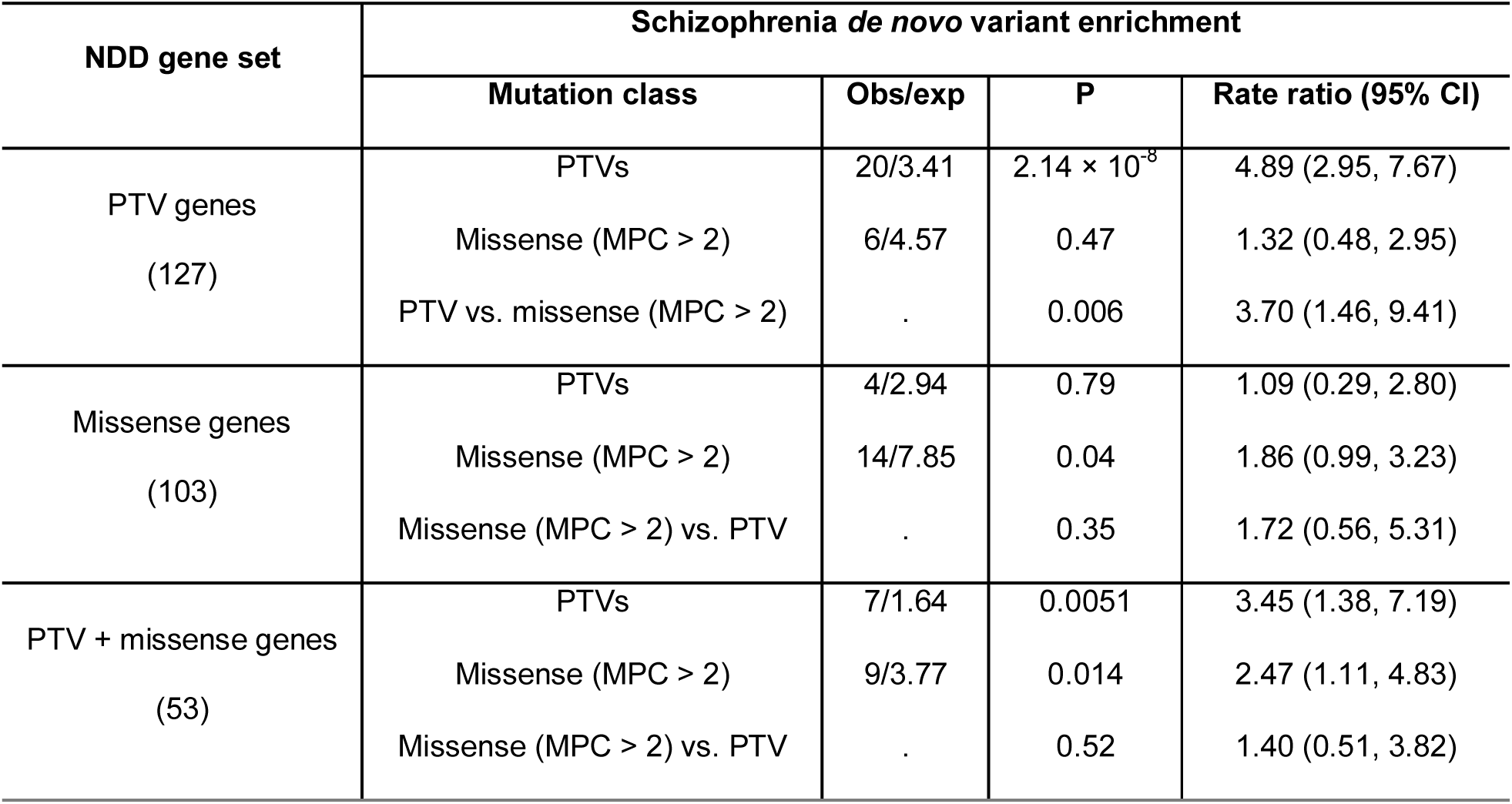
Enrichment of schizophrenia *de novo* variants in neurodevelopmental disorder (NDD) associated genes. Genes associated with PTV or missense *de novo* variants in NDDs (significance threshold set at P < 2.5 x 10^−6^) were evaluated for an enrichment of *de novo* variants in 3,444 schizophrenia trios using a two-sample Poisson rate ratio test. P-values are uncorrected and two-tailed. PTV/missense gene sets are defined as genes only associated with the given mutation class (i.e. excluding genes significant for the alternative mutation class) in the Deciphering Developmental Disorders study^13^. A Poisson regression model was used to evaluate differences between the rate of schizophrenia *de novo* PTVs and missense variants in NDD associated genes.

For 103 missense NDD genes, we again observed enrichment of the same class of *de novo* variant in schizophrenia (rate ratio = 1.86) and no enrichment for *de novo* PTVs (Table 1). The excess of schizophrenia *de novo* missense variants in NDD missense genes was greater than that observed for schizophrenia *de novo* PTVs (Table 1), but this was not statistically significant, possibly due to the small number of observations. For 53 genes that are independently associated with *de novo* PTVs and missense variants in NDDs, both classes of variant were enriched in schizophrenia (Table 1), thus providing further evidence that congruent classes of variants within genes confer risk for these disorders.

Across the full exome, we found that association test statistics per gene for PTV enrichment in schizophrenia was positively associated with the association test statistics for PTVs in NDDs (beta = 0.18; P = 2.97 x 10^−11^) but no evidence that PTV enrichment in schizophrenia was related to NDD missense test statistics (beta = −0.018; P = 0.67). We found a trend for missense enrichment in schizophrenia to be positively associated with the NDD missense test statistics (beta = 0.056; P = 0.069) but not with NDD PTV significance (beta = 0.033; P = 0.37). These results provide support for congruent variant classes, particularly for PTVs, contributing to risk for schizophrenia and NDDs beyond genes that meet exome-wide significance in NDD.

### Allelic pleiotropy

Of 46,772 unique single-nucleotide variants observed as *de novo* mutations in NDD studies (defined as NDD variants), 17 were also observed *de novo* in 3,444 schizophrenia trios (Table 2).

**Table 2.**
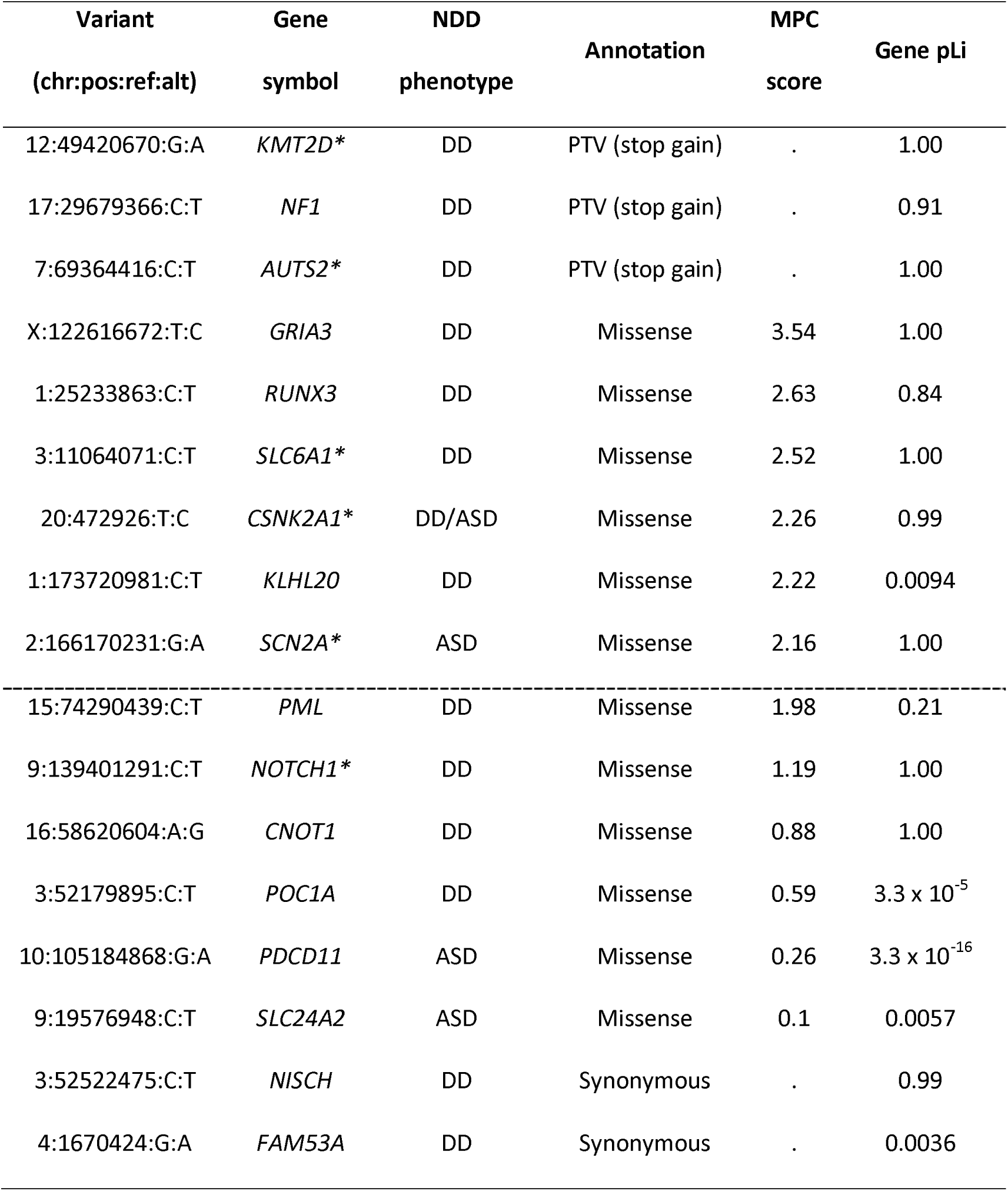
List of *de novo* variants observed in both schizophrenia and neurodevelopmental disorder trios. ‘NDD phenotype’ indicates whether the variant was reported in a developmental disorder (DD) trio or an autism spectrum disorder (ASD) trio^13,20^. Asterisks indicate genes associated at exome-wide significance with DNVs in NDDs. Variants above the dashed line were included in our primary test set. Variants below the line were included in our negative control set. Chr = chromosome; pos = genomic position (in build 37); ref = reference allele; alt = alternative allele; MPC = “Missense badness, Polyphen-2, Constraint” pathogenicity score^21^; pLi = “probability of loss-of-function intolerance”^22^.

| Variant<br>(chr:pos:ref:alt) | Gene<br>symbol | NDD<br>phenotype | Annotation | MPC<br>score | Gene pLi |
| --- | --- | --- | --- | --- | --- |
| 12:49420670:G:A | *KMT2D* * | DD | PTV (stop gain) | . | 1.00 |
| 17:29679366:C:T | *NF1* | DD | PTV (stop gain) | . | 0.91 |
| 7:69364416:C:T | *AUTS2* * | DD | PTV (stop gain) | . | 1.00 |
| X:122616672:T:C | *GRIA3* | DD | Missense | 3.54 | 1.00 |
| 1:25233863:C:T | *RUNX3* | DD | Missense | 2.63 | 0.84 |
| 3:11064071:C:T | *SLC6A1* * | DD | Missense | 2.52 | 1.00 |
| 20:472926:T:C | *CSNK2A1* * | DD/ASD | Missense | 2.26 | 0.99 |
| 1:173720981:C:T | *KLHL20* | DD | Missense | 2.22 | 0.0094 |
| 2:166170231:G:A | *SCN2A* * | ASD | Missense | 2.16 | 1.00 |
| 15:74290439:C:T | *PML* | DD | Missense | 1.98 | 0.21 |
| 9:139401291:C:T | *NOTCH1* * | DD | Missense | 1.19 | 1.00 |
| 16:58620604:A:G | *CNOT1* | DD | Missense | 0.88 | 1.00 |
| 3:52179895:C:T | *POC1A* | DD | Missense | 0.59 | $3.3 \times 10^{-5}$ |
| 10:105184868:G:A | *PDCD11* | ASD | Missense | 0.26 | $3.3 \times 10^{-16}$ |
| 9:19576948:C:T | *SLC24A2* | ASD | Missense | 0.1 | 0.0057 |
| 3:52522475:C:T | *NISCH* | DD | Synonymous | . | 0.99 |
| 4:1670424:G:A | *FAM53A* | DD | Synonymous | . | 0.0036 |

Schizophrenia *de novo* variants were significantly enriched for variants in the NDD primary set (9 observed, 1.20 expected; P = 5.0 x 10^−6^, Table 3). The enrichment for *de novo* primary NDD variants in schizophrenia was significantly greater than the general enrichment for the same types of *de novo* mutation in constrained genes and coding sequences (i.e. PTVs in LoF intolerant genes, and MPC ≥ 2 mutations) (P = 1.01 x 10^−5^; rate ratio (95% CI) = 6.91 (3.11, 13.38); Supplementary Table S1). The enrichment for specific NDD variants in schizophrenia is therefore not simply a reflection of the known modest excess of constrained *de novo* variants in the disorder. Schizophrenia *de novo* variants were not enriched in the negative control set (Table 3), which suggests that our finding is not caused by inaccuracies in estimating the expected *de novo* mutation rates.

**Table 3.** Enrichment of neurodevelopmental disorder variants in schizophrenia. The number of observed and expected *de novo* neurodevelopmental variants in 3,444 schizophrenia trios is shown. The primary variant set includes neurodevelopmental variants annotated as PTVs in genes with “probability of loss of function intolerance” (pLi) scores ≥ 0.9 or missense variants with MPC scores ≥ 2. The negative control variant set includes neurodevelopmental variants annotated as PTVs in genes with pLi scores < 0.9, missense variants with MPC scores < 2 and all synonymous variants. Enrichment statistics were generated using a Poisson rate ratio test. P-values are uncorrected and two-tailed. PTVs = Protein-truncating variants; NDD = neurodevelopmental disorders; CI = confidence interval.

| NDD variant set<br>(N variants) | Schizophrenia *de novo* variant enrichment |  |  |
| --- | --- | --- | --- |
|  | Observed / expected | P | Rate Ratio<br>(95% CI) |
| Primary<br>(5,863) | 9 / 1.20 | $5.0 \times 10^{-6}$ | 7.48 (3.42, 14.20) |
| Negative control<br>(40,909) | 8 / 9.86 | 0.75 | 0.81 (0.35, 1.60) |

We next looked at the specific classes of NDD mutation enriched in schizophrenia and found significant enrichment for both NDD PTVs (*P* = 0.01; rate ratio (95% CI) = 6.77 (1.40, 19.80)) and NDD missense variants (*P* = 0.00014; rate ratio (95% CI) = 7.90 (2.90, 17.19)) in the primary variant set (full details presented in Supplementary Table S2).

We sought replication of association between NDD variants in the primary set and schizophrenia using exome sequencing data from 4,070 schizophrenia cases and 5,712 controls. The rate of variants from the primary set was ∼2 fold higher in schizophrenia cases than in controls (P = 0.036; Table 4). The rate of both missense and PTV variants in the primary set were increased in schizophrenia cases compared with controls, although only the later was significantly higher (P = 0.024; Supplementary Table S3). The rate of NDD variants in the negative control set did not differ between cases and controls (Table 4). All NDD variants from the primary set observed in schizophrenia cases or controls are presented in Supplementary Table S4.

**Table 4.** Schizophrenia case-control analysis of neurodevelopmental disorders variants. Firth’s penalised logistic regression models were used to evaluate the burden of NDD variants in 4,070 schizophrenia cases and 5,712 controls. As this analysis included frameshift variants, the number of NDD variants in the primary and negative control set differs to that presented in Table 3. P-values are uncorrected and one-tailed. NDD = neurodevelopmental disorders; CI = confidence interval.

| NDD variant set<br>(N variants) | Schizophrenia case-control analysis |  |  |  |
| --- | --- | --- | --- | --- |
|  | Case rate<br>(N variants) | Control rate<br>(N variants) | P | Odds ratio (95% CI) |
| Primary<br>(7,979) | 0.0044<br>(18) | 0.0023<br>(13) | 0.036 | 1.90 (0.94, 3.95) |
| Negative control<br>(47,630) | 0.095<br>(389) | 0.097<br>(553) | 0.42 | 1.01 (0.89, 1.15) |

We were able to obtain additional phenotype data for each of the 9 schizophrenia probands who carried a *de novo* variant from the primary variant set (Supplementary Table S5). Four probands had an age of onset before 18. Two had evidence for lower than average premorbid cognitive function; one of those individuals had a *KMTD2* PTV, achieved normal developmental milestones, attended secondary school but left without qualifications and required supported employment, the other had an *AUTS2* PTV, delayed developmental milestones and required supported schooling for learning disability. None of the of other probands were reported to have developmental delay, intellectual disability, co-morbid ASD or pervasive developmental disorder.

Three of the 9 schizophrenia *de novo* variants in the primary variant set are recorded in the ClinVar database^23^ as pathogenic for Okur-Chung neurodevelopmental syndrome, Kabuki syndrome, and Neurofibromatosis type 1, which are all severe NDDs associated with intellectual disability (Supplementary Table S5). Four of the 9 schizophrenia *de novo* variants in the primary variant set are also recorded as pathogenic or likely pathogenic in the Decipher database^24^ (these include all three variants recorded as pathogenic in ClinVar). The remaining 5 variants are not present in either ClinVar or Decipher databases. Where available, the phenotypes of Decipher Patients carrying one of these pathogenic/likely pathogenic variants are presented in Supplementary Table S6; all Decipher patients had moderate/severe intellectual disability and/or developmental delay, among other neurodevelopmental phenotypes. A summary of the functions and conditions associated with genes affected by the primary variants observed as schizophrenia *de novo* variants is provided in Supplementary Table S7.

## Discussion

Previous comparisons of genetic liability based on rare CNVs and ultra-rare coding variants have shown evidence for genic pleiotropy between schizophrenia and NDDs^5,7–9^. Here, we extend that work to show that coding *de novo* variants in people with schizophrenia are most commonly (Table 1) of the same functional category as those that confer risk for NDDs. Moreover, we show that the specific *de novo* variants in people with NDDs are enriched in people with schizophrenia.

Given that neurodevelopmental impairment is typically more severe in NDDs than in schizophrenia^19^, we hypothesised that there would be a tendency for pleiotropic genes to be enriched for a more severe class of mutation in NDDs than in schizophrenia. However, in contrast to our expectation, we found that genes associated with *de novo* variants in NDDs were enriched for the same class of variant in schizophrenia, with the evidence being particularly strong for PTVs. Conversely, there was no evidence of schizophrenia *de novo* variant enrichment in genes associated with a different class of variant in NDD. We next conducted a more stringent analysis of allelic pleiotropy, by testing only the same set of rare coding variants in schizophrenia that occurred in people with NDDs. Here, our findings supported our hypothesis that constrained variants observed in NDDs are enriched in schizophrenia, thus providing evidence for pleiotropy at the allelic level.

Our study suggests that the same rare variants can confer risk to a range of neurodevelopmental and psychiatric outcomes, including DD, ASD and schizophrenia, thus supporting the hypothesis that at least some fraction of schizophrenia can be conceived of as part of a continuum of NDDs^19^. We did not find support for our prediction that the type of mutation is a major factor in determining the severity of neurodevelopmental outcome, though we cannot exclude the possibility that instances of this will become apparent through large-scale sequencing studies. Rather our findings suggest that clinical outcome reflects additional genetic, environmental or stochastic factors that can modify the effects of deleterious mutations. Indeed, there is evidence that the outcome for pathogenic CNVs is influenced by common genetic variation^25–27^. The situation for rare coding variants in schizophrenia is less clear^8^, but it is likely that similar considerations will apply. Our findings also indicate that the same changes in gene function can underlie both NDDs and schizophrenia, pointing to a shared molecular aetiology and therefore likely overlapping pathophysiology.

Schizophrenia and intellectual disability co-occur more often than is expected by chance, with around 3-5% of cases of schizophrenia being co-morbid^19,28^. The present findings further support the idea that co-morbidity is due to partly shared pathophysiology. However, it is unlikely that shared pathophysiology is restricted to those with co-morbid diagnoses. Many of the schizophrenia trios, including 3 of the 9 probands with a primary *de novo* NDD variant, were from studies that specifically excluded individuals with intellectual disability (Supplementary Table S8). All trios, apart from 17 trios taken from the Ambalavanan *et al* 2016 study^29^, had been recruited from general adult psychiatric services and all probands had received a primary DSM-IV or ICD-10 diagnosis of schizophrenia or schizoaffective disorder. Furthermore, only 2 of the 9 probands had evidence of clinically significant premorbid intellectual impairment, consistent with findings from other studies showing damaging *de novo* mutations may be enriched in, but are not confined to, those with intellectual disability^6,30^. These findings, together with the absence of evidence for pre-existing autism or related disorders (Supplementary Table S5), are of clinical relevance since they suggest that patients presenting with schizophrenia in the absence of neurodevelopmental comorbidities may carry damaging mutations that are associated with more severe neurodevelopmental outcomes. We also observed pleiotropic effects for variants known to be pathogenic for several syndromic developmental disorders, suggesting that schizophrenia should be included among the phenotypes associated with these mutations.

Given the substantial enrichment over expectation of primary NDD variants in our set-based analysis (∼7.5 fold) our analysis of allelic pleiotropy implicates variants in *KMT2D, NF1, AUTS2, GRIA3, RUNX3, SLC6A1, CSNK2A1, KLHL20*, and *SCN2A* in schizophrenia. However, definitive implication of any individual gene, much less any individual variant, requires much larger datasets than those currently available. Nevertheless, the following 4 variants are defined as pathogenic/likely pathogenic in the ClinVar^23^ and/or Decipher^24^ databases, which adds to the probability that they confer risk to schizophrenia: **1)** The missense variant in *CSNK2A1* (20:472926:T:C), which encodes an alpha subunit of casein kinase II, is pathogenic for Okur-Chung neurodevelopmental syndrome (OMIM #617062), an autosomal dominant disorder characterised by intellectual disability and dysmorphic facial features. **2)** The stop-gain PTV in *KMT2D*, which encodes a histone methyltransferase that methylates the Lys-4 position of histone H3, is pathogenic for Kabuki syndrome (OMIM#147920), an autosomal dominant disorder characterised by intellectual disability and additional dysmorphic facial features. **3)** The stop-gain PTV in *NF1* encodes neurofibromin and is pathogenic for neurofibromatosis, type 1, an autosomal dominant condition associated with learning disabilities and tumours of nerves and skin. **4)** The stop-gain PTV in *AUTS2* (Activator Of Transcription And Developmental Regulator) is recorded as likely pathogenic in Decipher. A summary of the 9 genes implicated in schizophrenia by our allelic analysis is presented in Supplementary Table S7.

In conclusion, we performed the first genome-wide study of allelic pleiotropy from rare coding variants for schizophrenia and NDDs. We show sets of genes associated with NDDs are enriched for congruent classes of variant in schizophrenia, and identify specific variants enriched for pleiotropic effects across both disorders. Collectively, our findings support the hypothesis that schizophrenia forms part of a continuum of NDDs including ASD and developmental disorders. Our study points to a shared molecular aetiology and the need for more work exploring the mechanistic and clinical relationships between NDDs and schizophrenia.

## Online methods

### Ethics statement

All research conducted as part of this study was approved by the Research Ethics Committee for Wales and consistent with regulatory and ethical guidelines.

### Schizophrenia *de novo* data

*De novo* variants from 3,444 schizophrenia proband-parent trios (2,121 male and 1,323 female probands) were obtained from 11 published studies (Supplementary Table S8)^6– 8,29,31–37^. The probands were ascertained from psychiatric wards or outpatient clinics, and all had received a DSM-IV (Diagnostic and Statistical Manual of Mental Disorders; fourth edition) or ICD-10 (International Statistical Classification of Diseases and Related Health Problems; 10th revision) research diagnosis of schizophrenia or schizoaffective disorder, apart from 5 probands who had a diagnosis of non-organic psychosis (details in Supplementary Table S8).

*De novo* variants were re-annotated using Ensemble Variant Effect Predictor (version 96)^38^. PTVs included stop-gain, frameshift, or splice donor/acceptor variants. Missense variants were annotated with their “Missense badness, Polyphen-2, constraint” (MPC) score, which is a pathogenicity metric that combines predictions of variant deleteriousness with measures of regional missense constraint^21^. We prioritised missense variants with MPC scores ≥ 2 in our analyses, as this class of variant has been shown to be enriched in ASD cases compared with controls^20^.

### Genic pleiotropy

#### Neurodevelopmental disorder gene sets

NDD associated genes were identified from the Deciphering Developmental Disorders study^13^. In that study, 180 and 156 genes were, respectively, associated with *de novo* PTV and missense variants at exome-wide significance (P value < 2.5 x 10^−6^). 53 genes were independently associated at this threshold with both PTVs and missense variants. We stratified the NDD associated genes into 3 independent groups – PTV specific (127 genes), missense specific (156 genes) and PTV + missense (53 genes) – and tested each group for enrichment for *de novo* variants in the schizophrenia probands. The genes included in these sets are provided in Supplementary Table S9. We did not include ASD associated genes in these sets as independent PTV and missense P values were not reported in the largest published ASD study^20^.

#### Statistics

For 3,444 schizophrenia trios, we used published gene mutation rates to estimate the number of *de novo* variants expected to occur under the null in the NDD gene sets^39,40^. Where possible, gene mutation rates were adjusted for sequencing coverage; the use of unadjusted per-gene mutation would overestimate the expected number of *de novo* variants in these trios, and produce more conservative enrichment results (see^8^ for further details). A two-sample Poisson rate ratio test was used to compare the enrichment of *de novo* variants in NDD genes, relative to the expected number, with the enrichment observed for all genes outside of the NDD gene set relative to the expected number, thereby controlling for the minimal elevation in the background schizophrenia *de novo* rates. Gene set enrichment tests were conducted for two mutation classes: PTVs and missense variants with MPC scores ≥ 2.

A Poisson regression was used to test for differences in the degree of enrichments for schizophrenia PTV and missense *de novo* variants in NDD gene sets. Here, a regression was first performed for each mutation class, with the number of observed *de novo* variants being the outcome variable, gene set membership (e.g. NDD associated or not) a categorical predictor, and the log of the expected number of *de novo* variants in each gene set category the offset. The log of the rate ratio for the enrichment of NDD associated genes in PTVs relative to missense variants is then the difference in the log rate ratios for NDD genes in the two Poisson regressions (i.e. the regression coefficient for NDD gene membership). The variance of this difference is the sum of the variances of the regression coefficients, enabling confidence intervals to be generated. The square of the difference in regression coefficients divided by the sum of the variances can be compared to a χ^2^ distribution with one degree of freedom to give a test of significant differences in the schizophrenia enrichment of PTV and missense *de novo* variants in NDD associated genes. This approach allows for the background enrichment of schizophrenia *de novo* variants in non-NDD genes to differ between PTV and missense variants.

We also used a Poisson regression model to evaluate the relationship between schizophrenia *de novo* variant enrichment and gene level P values for PTV and missense variants in NDDs simultaneously. NDD gene P values were taken from^13^. Unlike the gene set analysis, which required an arbitrary significance threshold for a gene being considered NDD associated (i.e. P < 2.5 x 10^−6^), this Poisson regression was applied to all genes.

N SZ variants (per gene) ∼ -log(DD PTV P value) + -log(DD missense P value), offset(log(N SZ expected variants))

### Allelic pleiotropy

#### Neurodevelopmental disorder variants

NDD variants were identified from *de novo* variants observed in the largest ASD and DD proband-parent sequencing studies (total NDD trios = 37,488; Table 5), which together reported a total of 48,155 single-nucleotide *de novo* variants, corresponding to 46,772 unique single-nucleotide variants (summarised in Table 5, full list of variants in Supplementary Table S10). We divided these variants into primary and negative control sets. The primary set contains variants with characteristics known to be associated with pathogenicity for NDDs^13,41^, namely PTVs in loss-of-function intolerant genes (genes with gnomAD pLi scores ≥ 0.9^22^) and missense variants with MPC scores ≥ 2^21^. The negative control set contained all remaining variants (PTVs in genes with pLi scores < 0.9, missense variants with MPC scores < 2 and all synonymous variants), properties that do not predict NDD pathogenicity. Under an allelic pleiotropy model, we predicted that schizophrenia *de novo* variants would be more enriched among the primary variant set than the negative control variant set.

**Table 5.** Summary of single-nucleotide variants included in the allelic pleiotropy *de novo* analysis. The ‘N DNVs’ column shows the total number of *de novo* missense, synonymous, stop-gain, splice-donor or splice-acceptor variants reported in the respective phenotype after excluding variants on the Y chromosome or in mitochondrial DNA. The Unique DNVs column shows the number of *de novo* variants observed in the respective phenotype after excluding duplicate variants. DNV = *de novo* variant; PTV = protein truncating variant; miss = missense variant, syn = synonymous variant.

| Phenotype | N trios | Total DNVs<br>(PTVs, miss, syn) | Unique DNVs<br>(PTVs, miss, syn) |
| --- | --- | --- | --- |
| Schizophrenia | 3,444 | 3,208<br>(186, 2197, 285) | 3,207<br>(186, 2197, 824) |
| DD<sup>13</sup> | 31,058 | 40,818<br>(3,638, 28,193, 8,987) | 39,560<br>(3,400, 27,211, 8,949) |
| ASD<sup>20</sup> | 6,430 | 7,337<br>(516, 4,954, 1,867) | 7,306<br>(514, 4,934, 1,858) |
| NDD (ASD + DD) | 37,488 | 48,155<br>(4,154, 33,147, 10,854) | 46,772<br>(3,900, 32,076, 10,796) |

#### Statistics

Tri-nucleotide mutation rates were used to estimate the expected per-generation mutation rates for NDD variants^21^. These mutation rates were then used to derive the number of NDD variants expected to occur *de novo* under the null hypothesis in the 3,444 schizophrenia trios. As mutation rates have not been empirically established for indels, only single-nucleotide variants were considered (Table 5).

The numbers of schizophrenia *de novo* variants overlapping our primary and negative control variant sets were compared to that expected under the null using a two-tailed Poisson exact test. We also used a two-sample Poisson exact test to evaluate whether the enrichment of schizophrenia *de novo* variants in the primary variant set was greater than the schizophrenia background *de novo* rate of all PTVs in LoF intolerant genes and missense variants with MPC scores ≥ 2. Statistics were generated using R statistical software (version 3.4.3) and the poisson.test() function.

NDD variants in our primary and negative control sets were further evaluated using a Swedish schizophrenia case-control exome sequencing data set, which consists of 4,079 cases and 5,712 controls^9^. Case-control exome sequencing data were analysed using Hail (https://github.com/hailis/hail). To test for an excess burden of NDD variants in cases compared with controls, a one-tailed Firth’s penalized-likelihood logistic regression model was used, correcting for the first 10 principal components derived from the sequencing data, and for the exome-wide burden of synonymous variants, sequencing platform and sex. To focus the case-control analysis on ultra-rare alleles, as those are more likely to be pathogenic, we excluded variants with an allele count > 5 in gnomAD^22^. Frameshift variants were included in the case-control analysis.

## Acknowledgements

The work at Cardiff University was supported by Medical Research Council Centre Grant No. MR/L010305/1 (to MJO) and Program Grant No. G0800509 (to MJO, MCO, JTRW, PH).

## DECIPHER Acknowledgments

This study makes use of data generated by the DECIPHER community. A full list of centres who contributed to the generation of the data is available from https://decipher.sanger.ac.uk and via email from. Funding for the DECIPHER project was provided by Wellcome. We acknowledge Drs. Peter Turnpenny, Fernando Santos Simarro, Ramsay Bowden, Joanna Jarvis, Jenny Carmichael and Andrew Green for providing their consent to publish the phenotypes observed in DECIPHER patients who have a primary neurodevelopmental disorder variant that is also observed in a schizophrenia patient. We acknowledge that those who carried out the original analysis and collection of the DECIPHER data bear no responsibility for the further analysis or interpretation of the data.

## Swedish Exome Sequencing Acknowledgments

The datasets used for the analysis described in this manuscript were obtained from dbGaP at http://www.ncbi.nlm.nih.gov/gap through dbGaP accession number phs000473.v2.p2. Samples used for data analysis were provided by the Swedish Cohort Collection supported by the NIMH Grant No. R01MH077139, the Sylvan C. Herman Foundation, the Stanley Medical Research Institute and The Swedish Research Council (Grant Nos. 2009-4959 and 2011-4659). Support for the exome sequencing was provided by the NIMH Grand Opportunity Grant No. RCMH089905, the Sylvan C. Herman Foundation, a grant from the Stanley Medical Research Institute and multiple gifts to the Stanley Center for Psychiatric Research at the Broad Institute of MIT and Harvard.

## Author contributions

ER, PH, MJO and MCOD conceived and designed the research. ER analysed the data. ER, PH, JTRW, MJO, MCOD, HDJC and RR contributed to the interpretation of the results. HH, WJC, MT, SJG, GK, MCOD and MJO led the acquisition of the clinical samples. ER, MJO, MCOD wrote the manuscript, which was read, edited and approved by all authors.

## Data availability

All schizophrenia *de novo* variants were obtained from the published sources outlined in Supplementary Table S8. *De novo* variants from ASD trios were obtained from Satterstrom et al 2020^20^, and *de novo* variants from DD trios were obtained from Kaplanis et al 2019^13^. DD gene level association statistics were obtained from Kaplanis et al 2019^13^.

