## Supplementary Table S7 for "Schizophrenia, autism spectrum disorders and developmental disorders share specific disruptive coding mutations"

**Supplementary Table S7**. Functions and conditions associated with genes affected by the primary neurodevelopmental disorder variants observed as schizophrenia de novo variants.

| **Gene ID** | **Gene Name** | ***Alternative ID*** | **General Function** | **Related Conditions** | **Expression Profile** | **Key References** | **Notes** |
| --- | --- | --- | --- | --- | --- | --- | --- |
| *KMT2D* | Lysine methyltransferase 2D | *ALR, MLL2, MLL4* | A methyltransferase, responsible for a majority of the H3K4me1/2 marks in mammalian cells along with KMT2C.  Together they have been linked to a role in neuronal differentiation.  KMT2D plays critical roles in regulating development, differentiation, metabolism, and tumour suppression. | Kabuki syndrome ( Autosomal Dominant) | Ubiquitous expression across most bodily tissues, including brain (RPKM 3.2).  Important for embryonic development. | ^1–9^ | The encoded protein is part of a large protein complex called ASCOM.  ASCOM is a transcriptional regulator of the beta-globin and oestrogen receptor genes  Deficiency compromises the development of regulatory T cells (T-reg cells).  **N.B.** The similar gene KMT2F (also known as SET1, SETD1A) linked to SZ.  **N.B.B.** *KMT2C* has been implicated in Kleefstra syndrome.  ***N.B.B.B.*** *Kmt2a/Mixed-lineage leukemia 1* (*Mll1*), in mouse postnatal forebrain and adult prefrontal cortex (PFC) is associated with increased anxiety and cognitive deficits. |
| *NF1* | Neurofibromin 1 | *NFNS, VRNF, WSS* | Functions as a negative regulator of the Ras signal transduction pathway by stimulating GTPase activity of Ras. | Neurofibromatosis 1 – characterised by tumours of nerves and skin (neurofibromas),  Learning disabilities  Autosomal Dominant | Ubiquitous expression across most bodily tissues, highly expressed in brain (RPKM 8.2) and thyroid (RPKM 9.3).  **N.B.** Homozygous LOF mutation unviable in mice. | ^10–14^ | Over 50% of individuals with NF1 experience learning disabilities. |
| *AUTS2* | Activator of transcription and developmental regulator | *FBRSL2, MRD26* | Component of a Polycomb group (PcG) multiprotein PRC1-like complex.  The PcG PRC1 complex mediates transcriptional activation of CNS genes via epigenetic remodelling of histones and chromatin.  AUTS2 regulates neuritogenesis via activation of Rac1 signalling and is involved in neural migration during embryo development. | Autism Spectrum Disorder (ASD), Schizophrenia  Autosomal Dominant | Ubiquitous expression across most bodily tissues including brain (RPKM: 1.95) brain.  **N.B.** Homozygous LOF mutation unviable in mice. | ^15–20^ | Implicated in neurodevelopment and as a candidate gene for numerous neurological disorders, including autism spectrum disorders, intellectual disability, and developmental delay. |
| *GRIA3* | Glutamate Ionotropic Receptor AMPA Type Subunit 3 | *GLUR3, GLURC, MRX94* | Glutamate receptor that functions as ligand-gated ion channel in the central nervous system. It plays an important role in excitatory synaptic transmission.  Codes for AMPA receptor subunit.  Mutations result in absent or diminished GluA3 protein. | Mental retardation (X-linked, Syndromic, wu type),  Intellectual Disability,  ASD, Major Depressive Disorder (sleep disturbances),  Methamphetamine Dependence (& associated psychosis)  Deletion or duplication can cause ID. | Specific expression in brain (RPKM: 19.9), adrenal: (RPKM 3.0).  Important in embryonic brain development. | ^21–27^ | These receptors are heteromeric protein complexes composed of multiple subunits, arranged to form ligand-gated ion channels and are activated in a variety of normal neurophysiologic processes.  Increased levels of dopamine in the striatum of *Gria3KO* mice  **N.B.** Previously considered a BPD candidate gene. |
| *RUNX3* | Runt-related transcription factor 3 | *AML2 , CBFA3, PEBP2aC* | Forms the heterodimeric complex core-binding factor (CBF) with CBFB.  Regulator of CD8+ T-cell thymocyte development. | Cancer metastasis,  Pre-eclampsia,  Placental Dysfunction | Biased expression in bone marrow (RPKM: 19.5) and spleen (RPKM16.7)  **N.B.** Homozygous LOF mutants unviable in mice. | ^28,29^ | Under epigenetic regulation. |
| *SLC6A1* | Voltage-dependent c-aminobutyric acid (GABA) transporter 1 (GAT-1) protein | *GABATHG, GABATR, GAT1 MAE* | Terminates the action of GABA by its high affinity sodium-dependent reuptake into presynaptic terminals. | Schizophrenia, ADHD,  Epileptic encephalopathy | SLC6A1 is primarily expressed in the adult brain (RPKM: 26.6), specifically, in GABAergic neurons and astrocytes.  Essential for embryonic development. | ^30–34^ | Mutations often result in loss of function of GAT-1 and thus reduced GABA re-uptake from the synapse. |
| *CSNK2A1* | Casein Kinase 2 Alpha 1 | *CK2A1, Cka1,*  *Cka2* | Casein kinase II is a serine/threonine protein kinase that phosphorylates acidic proteins such as casein. It is involved in various cellular processes, including cell cycle control, apoptosis, and circadian rhythm. | Neurodevelopmental disorders (Okur-Chung Neurodevelopmental Syndrome),  Intellectual disability,  Seizures | Ubiquitous expression across adult tissues including brain (RPKM: 25.8)  Essential for embryonic development. | ^35–38^ | Often associated dysmorphic facial features. Congenital heart abnormalities identified in nearly 30% of the patients with *CSNK2A1* mutations. |
| *KLHL20* | Kelch Like Family Member 20 | *KLEIP, KLHLX, KHLHX* | KLHL20 is a related CUL3-dependent ubiquitin ligase linked to autophagy, cancer, and Alzheimer's disease that promotes the ubiquitination and degradation of substrates including DAPK1, PML, and ULK1. | Alzheimer’s disease,  Cancer | Ubiquitous expression across adult tissues including brain (RPKM: 4.7).  Important in Embryonic development. | ^39–42^ | Stress response of KLHL20 is linked to neurodegeneration. KLHL20 RNA transcript levels being among the top 20 biomarkers for Alzheimer’s disease progression. |
| *SCN2A* | Sodium Voltage-Gated Channel Alpha Subunit 2 | *BFIC3, BFIS3, BFNIS* | Mediates the voltage-dependent sodium ion permeability of excitable membranes.  Implicated in the regulation of hippocampal replay occurring within sharp wave ripples (SPW-R) important for memory | Autism Spectrum Disorder (ASD) | Biased expression in brain (RPKM: 20.5). | ^43–45^ | Variants with these conditions are usually truncating, suggesting haploinsufficiency plays a major role. |

Key references from Supplementary Table S7
